# Physiological basis of noise-induced hearing loss in a tympanal ear

**DOI:** 10.1101/698670

**Authors:** Ben Warren, Georgina E Fenton, Elizabeth Klenschi, James FC Windmill, Andrew S French

## Abstract

Acoustic overexposure, such as listening to music too loud and too often, results in noise-induced hearing loss. The pathologies of this prevalent sensory disorder begin in the synapses of the primary auditory receptors, their postsynaptic partners and supporting cells. The extent of noise-induced damage, however, is determined by over-stimulation of primary auditory receptors. When over-stimulated, an excessive amount of positive ions flood into the primary auditory receptors, triggering the activation of ion channels and possibly disrupting their ability to encode sound. A systematic characterisation of the electrophysiological function of primary auditory receptors is warranted to understand how noise-exposure impacts on downstream targets, where the pathologies of hearing loss begin. Here, we used the experimentally-accessible locust ear to characterise noise-induced changes in the auditory receptors. Although, we found a decrease in ability of the primary auditory neurons to encode sound, this is probably due to pathologies of their supporting cells.

## Introduction: Consequences of noise-induced hearing loss across animal ears

Our senses endow us with an incredible subjective experience of our world. The gatekeepers of this external sensory environment – sensory receptors – selectively encode stimuli into electrical signals to give us our five senses. Overstimulation of sensory neurons, however, leads to damage and the sound-sensitive cells in our ears are particularly prone to permanent injury (Lundström & Johansson, 1986; Richard et al., 2008; Young, 1988). As a result, the most common sensory impairment in humans is hearing loss and the majority of preventable hearing loss is due to excessive noise exposure.

The ears of all animals have specialised ciliated sound-sensitive receptor cells, likely of a common evolutionary origin (Fritzsch & Beisel, 2004), that transduce sound into electrical potentials. Critical to this process of auditory transduction are stretch-sensitive ion channels, embedded in the membranes of the cilia, which open in response to sound (Joerg & Kozlov, 2016; Gillespie & Walker, 2001). Once open, cations flow into the auditory receptor cells to generate the primary electrical signal – the transduction potential. In vertebrates this transduction potential triggers the release of neurotransmitter in the base of their specialised non-spiking multi-ciliated auditory receptor cells – so-called hair cells (Moser et al., 2006; Fuchs, 2005). Postsynaptic neurons then convert this synaptic signal into spikes that travel to the central nervous system to give the percept of sound (Pickles, 1982). In insects, transduction and spike-generation take place within the same auditory neuron, which bears a solitary sensory cilium where the transduction ion channels are housed. In Orthoptera, such as locusts and crickets, the transduction potential is converted into a small dendritic spike that travels along a single dendrite of the neuron (Hill, 1983b; Oldfield & Hill, 1986) to the soma. There, most probably in the axon hillock, it triggers a larger axonal spike that carries auditory information to the central nervous system (Warren & Matheson, 2018).

Despite the different architecture of auditory receptors across the animal kingdom, they all suffer from excessive cation influx when acoustically overstimulated. In mammals, excessive calcium entry leads to production of reactive oxygen species by mitochondria (Böttger & Schacht, 2013; Fischel-Ghodsian et al., 2004) glutamatergic excitotoxicity (Kujawa & Liberman, 2009; Pujol et al., 1993), a decrease in synapses (Fernandez et al., 2015; Liberman and Liberman, 2015; Furnam et al., 2013; Lin et al., 2011), which can trigger apoptotic and necrotic pathways in the hair cells (Wagner & Shin, 2019). But excessive acoustic stimulation triggers pathologies, not only in the primary auditory receptors themselves, but in their postsynaptic partners (Sharon et al., 2009; Robertson, 1983), other supporting cells (Shi & Nuttall, 2003) and later auditory processing areas in the central nervous system (Fetoni et al., 2013; Noreña et al., 2010; Wei–ju et al., 2010). Similar pathologies manifest in invertebrate hearing models (Christie et al., 2013) but there is no systematic understanding of changes in the electrophysiological properties and function of the auditory receptor cells themselves shortly after noise exposure for any animal ear.

Here, we used the physiologically-accessible locust ear to measure electrophysiological changes in the auditory neurons directly after noise exposure, nerve potentials from the auditory nerve and mechanical changes in the vibrations of the tympanum. The locust ear permits a detailed characterisation of the sound-elicited ionic currents because whole-cell patch-clamp recordings can be conducted during concurrent stimulation by airborne sound in an intact ear (Warren and Matheson, 2018) - an approach not possible in other hearing models. In addition, composite spikes can be recorded from the auditory nerve and the tympanum is easily assessable for mechanical analysis (Windmill et al., 2005). The locust’s auditory Müller’s Organ is attached directly to the inside of the tympanum – like a splayed hand - and has three groups of neurons that attach onto discrete parts of the tympanum to exploit its heterogeneous resonant frequencies to detect sound from 200 Hz to 40 kHz (Jacobs et al., 1999). We focus our analyses on Group III auditory neurons that make up ~46 of the 80 auditory neurons and are broadly tuned to 3 kHz (Warren & Matheson, 2018). In response to acoustic overstimulation we measured physiological changes in the sound-evoked displacements of the tympanum, electrical responses from the auditory nerve, ionic currents in individual auditory neurons, and relative abundances of some putative sound-activated ion channel genes.

## Results

### Biomechanics of the tympanum

We first focused our analyses on *in vivo* measures of hearing impairment after auditory overexposure. To do this, Elizabeth Klenschi exploited the accessible nature of the locust ear to measure *in vivo* tone-evoked vibrations from the external surface of the tympanum, directly opposite where Group III auditory neurons are attached on the inside of the tympanum. For multiple auditory systems across phyla (including humans) the health of the auditory receptor cells can be surveyed from the sounds produced by mechanical vibrations of the tympanum (Kemp, 2002). Therefore, Elizabeth measured the displacement and mechanical gain of the tympanum. Mechanical gain is the amplitude of vibrations as a function of sound amplitude. The displacement of the tympanum and its mechanical gain was increased across SPLs from 50-100 dB SPL for noise-exposed locusts (Figure 1Ai, 2Aii).

**Figure 1.**
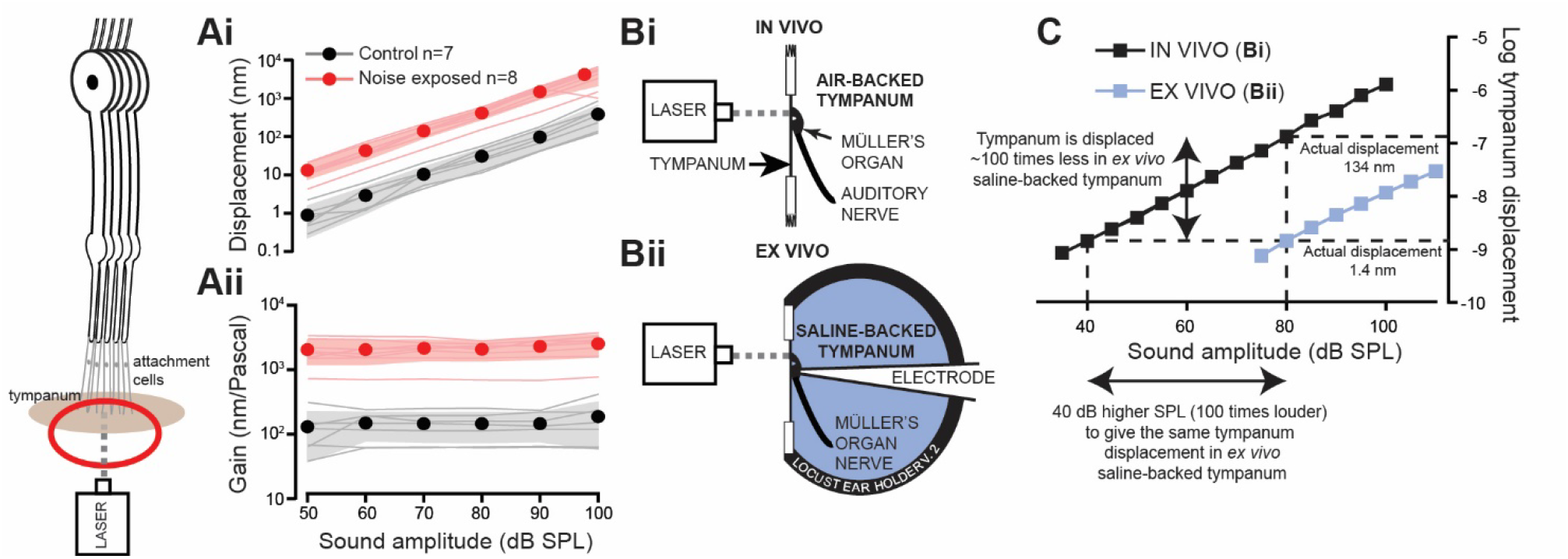
*In vivo* Doppler laser measurements of tympanal displacements shows increased displacements and gain for noise-exposed locusts. **Ai.** The displacement of the tympanum was higher for noise-exposed locusts (Linear Mixed Effects Model (LMEM) t=_(86.0)_4.829, p<0.0001, effect size d=0.75896). **Aii** The gain, nm displacement per Pascal, was also higher for noise-exposed locusts (LMEM t=_(79.18)_-3.432, p=0.0009, effect size d=3.44854) (Control: n=7, N=7; Noise-exposed: n=8, N=8). **Bi**. The experimental setup for *in vivo* recording from an intact ear. **Bii** Experimental setup for *ex vivo* recording with a saline-backed tympanum, necessary for intracellular recordings from individual auditory neurons. **C.** Comparison of tympanal displacements in an *in vivo* and *ex vivo* preparation backed by air and saline respectively. A 40 dB louder tone is required to move the tympanum by the same amount when backed by saline (*ex vivo*) compared to air (*in vivo*).

**Figure 2.**
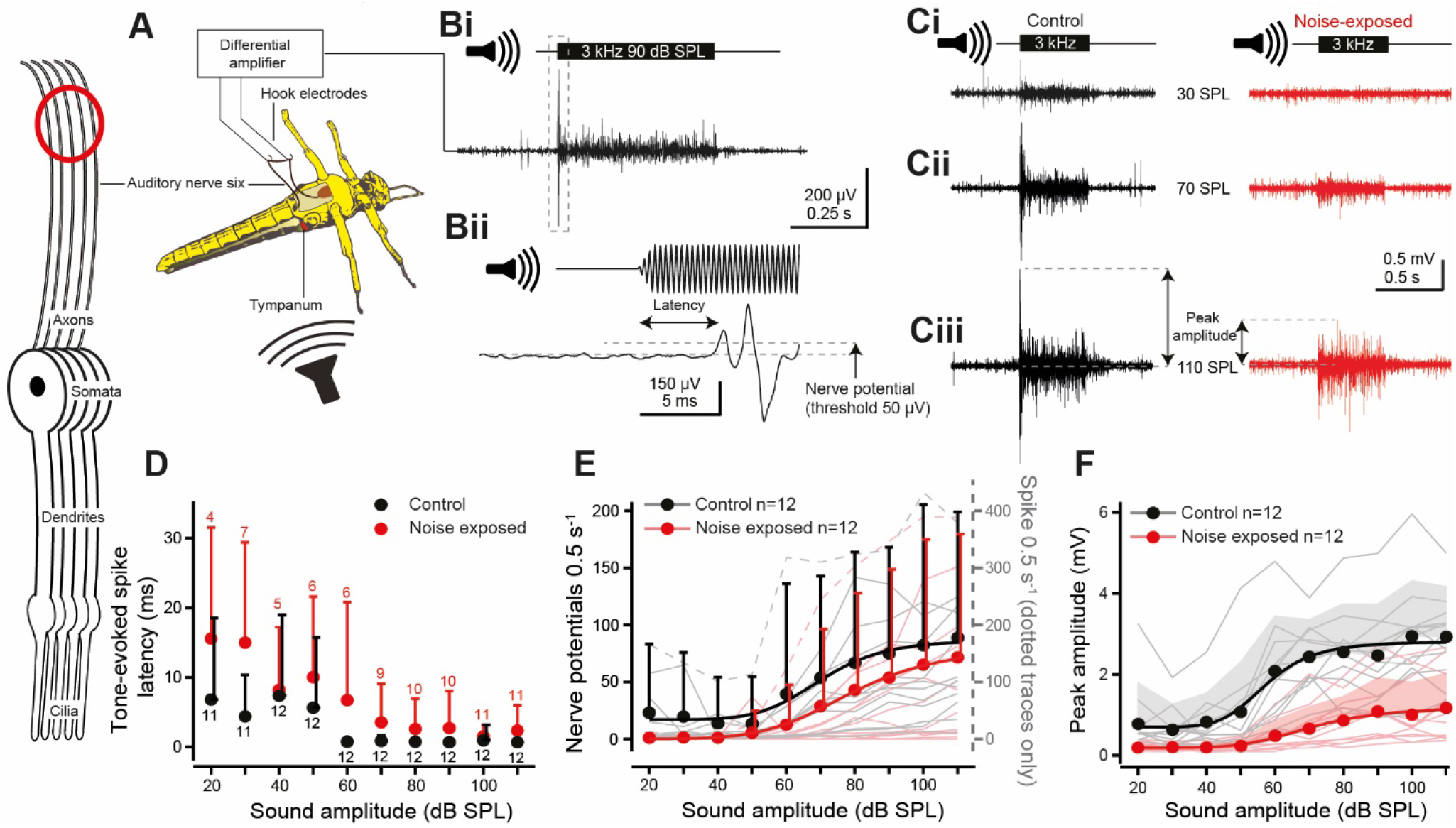
*In vivo* hook electrode recordings from the auditory nerve (six). Neuron schematic on left, with red circle, indicates where the electrical signals were recorded – from the auditory nerve. **A**. Schematic showing the vivisection and placements of the hook electrodes under the auditory nerve. **Bi.** Example recording using a 3 kHz tone at 90 dB SPL **Bii** with an expanded view (of dotted box in **Bi**) to show latency to first nerve potential and the 50 μV threshold used to count nerve potentials. **C.** Nerve responses for a 0.5 s 3 kHz tone at **Ci** 30, **Cii** 70, and **Ciii** 110 dB SPL (peak amplitude calculation for **F** is shown with double headed arrows). **D.** Quantification of the latency to first nerve potential in response to a 3 kHz tone for control and noise-exposed locusts. Noise-exposed locusts had delayed spike generation compared to their control counterparts (LMEM t=_(213.9)_0.457 p=0.001, effect size d=0.666374). Means are plotted as circles, positive standard deviation is plotted as error bars. **E**. Quantification of the number of tone-evoked nerve potentials above 50 μV, which increased for louder sound amplitudes (LMEM t_(694.0)_=9.717, p<0.0001, effect size=0.238436). The number of nerve potentials were not different between control and noise-exposed locusts (LMEM t=_(27.27)_0.690, p=0.496). Nerve potential counts are well fitted with Boltzmann equations (solid lines, Control R^2^=0.962; Noise-exposed R^2^=0.997). Means are plotted as circles, positive standard deviation is plotted as error bars (the error bars for noise-exposed means are offset, right, for figure clarity). Nerve potential counts from individual locusts are plotted as thin shaded lines. (N.B. Two individual particularly high neve potential counts, included in the analysis, are plotted as dotted lines on the right axis. **F.** The peak amplitude response of control locusts was higher than noise-exposed locusts at higher SPLs (LMEM: Treatment x SPL interaction, t_(649.0)_=10.00 p<0.0001, effect size d=1.138042). The peak response increased for higher sound amplitudes and was well fitted with a Boltzmann equation (solid lines, Control R^2^=0.973; Noise-exposed R^2^=0.986) (Control, n=12, N=12; Noise-exposed, n=12, N=12).

In order to quantitatively record tone-evoked currents and spikes from individual auditory neurons the ear was excised from the locust, placed in a recording chamber and the internal surface, with Müller’s organ, perfused with saline. Immersion of Müller’s organ in saline is necessary to gain access to, image and record from individual auditory neurons through whole-cell patch-clamp intracellular recordings (Warren & Matheson, 2018). Elizabeth quantified the reduction in tone-evoked tympanal vibrations when the tympanum is backed by water – as opposed to its tracheal air backing when *in vivo* (Figure 1Bi, Bii). Elizabeth found that vibrations of the tympanum were 100 times lower when the tympanum was backed by saline or, put another way; she required a tone 40 dB louder to move the tympanum by the same amount when backed by saline as opposed to air (Figure 1C).

### In vivo electrophysiology

Next, we investigated the effect of noise-exposure on the ability of auditory neurons to produce sound-evoked spikes. To accomplish this, Georgina Fenton recorded tone-evoked nerve potentials from the auditory nerve (nerve six) with hook electrodes in an *in vivo* preparation that left the first abdominal segment, with its bilateral ears, intact (Figure 2A). The nerve potentials are a read out of the population activity of Müller’s organ auditory neurons. The majority of the auditory neurons are broadly tuned to 3 kHz (Jacobs et al., 1999), so Georgina used a 3 kHz tone to elicit nerve potentials which were recorded as multiphasic extracellular potentials with a short delay (Figure 2Bi, ii). An increased sound amplitude elicited an increase in the nerve potentials (Figure 2Ci, ii, iii). Across hearing models hearing impairment is characterised by an increase in the latency of tone-evoked electrical signals (Christie & Eberl, 2014; Coomber et al., 2014; Van Heusden & Smoorenburg, 1981; Salvi et al., 1980).

Noise-exposed locusts replicated other hearing models with delayed spike generation in their auditory nerves compared to control locust ears (Figure 2D). Both control and noise-exposed locusts’ spike latency were reduced at higher sound amplitudes. Tone-evoked nerve potentials, above 50 μV in amplitude, were positively correlated with sound amplitude which was well fitted with a Boltzmann equation (Figure 2E, solid lines) but noise-exposure had no significant decrease on the number of nerve potentials. Georgina also measured the peak amplitude of the nerve potentials (Figure 2Ciii, double-headed arrows), which is the summated response of multiple spikes, typically elicited at tone onset. The peak amplitude was well fitted with a Boltzmann relationship, and the peak amplitude of noise-exposed locusts was below that of their control counterparts across SPLs (Figure 2F).

### Auditory neuron morphology, electrical properties and transduction current

Ben Warren measured auditory neuron morphology, characterised their electrical properties and measured their sound-elicited currents through whole-cell patch-clamp recordings from individual neurons from excised ears. Group III neurons in Müller’s organ (Figure 3A,B,C) have a long ~100 μm dendrite whose length and diameter was not different between control and noise-exposed locusts (Figure 4C, Table 1). In addition, there were no large differences in the membrane potential, membrane resistance and capacitance between noise-exposed and control auditory neurons (Table 1).

**Figure 3.**
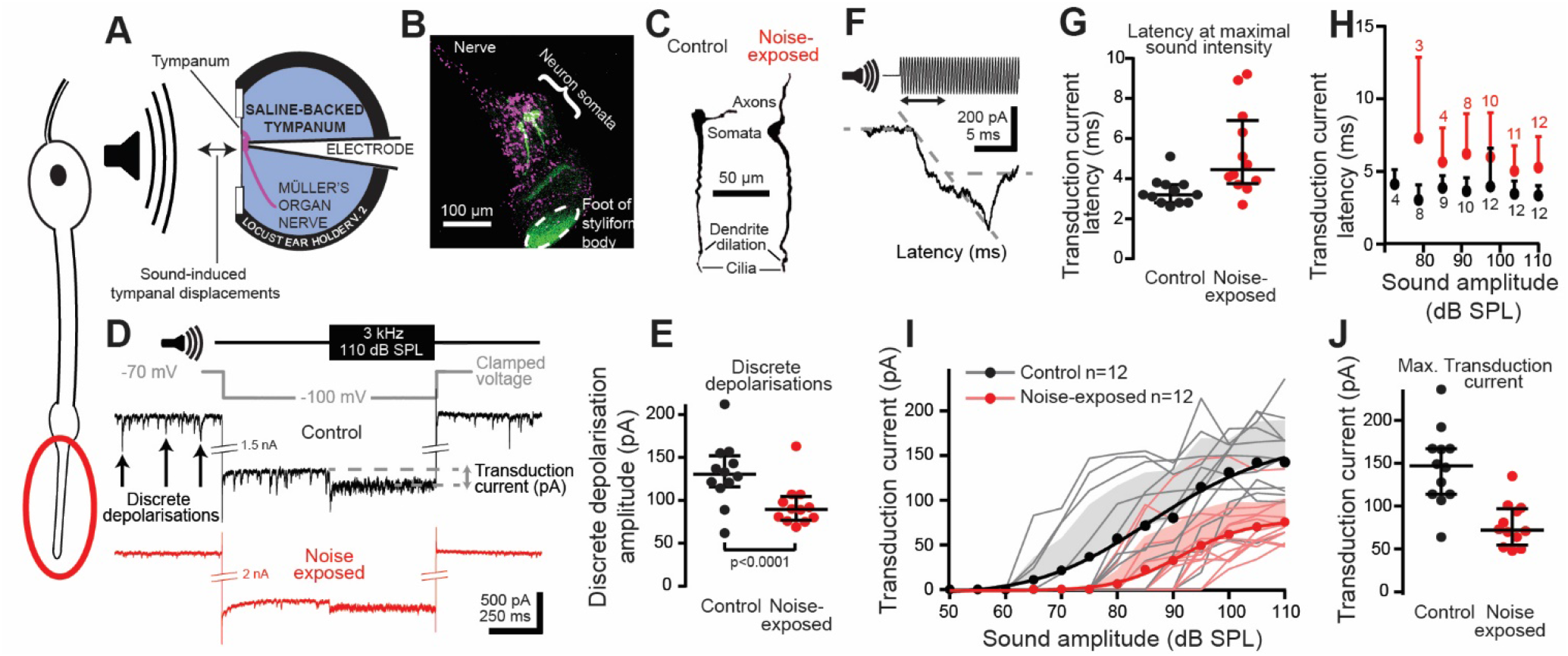
Intracellular whole-cell patch-clamp recordings of auditory neurons, their spontaneous and tone-evoked transduction ion channel activity and neuron morphology. All spontaneous and tone-evoked transduction currents occur in the auditory neurons’ cilium highlighted by a red circle on the neuron schematic on the left. **A.** Schematic showing experimental setup for intracellular patch-clamp recordings from Müller’s organ housed on the internal surface of the tympanum perfused with saline. The outside of the tympanum is acoustically driven by airborne sound by a speaker. **B.** Double staining of the nuclei of cells of Müller’s organ (magenta, DAPi) and three Group-III auditory neurons individually stained with neurobiotin/Dylight 488 streptavidin (green). There is some non-specific green fluorescence at the point of attachment of the organ to the tympanum (foot of styliform body). **C.** A neurobiotin-streptavidin staining of two Group-III auditory neurons *in situ* reveals the dendrite dilation and apical cilium. **D.** Sound stimulation, voltage-clamp and recording protocol used to maximise the transduction current during intracellular patch-clamp recordings. A 3 kHz tone at 110 dB SPL (top trace) was used to stimulate Group-III auditory neurons at their most sensitive frequency. The neurons were voltage-clamped to −100 mV (grey trace) to increase the electrochemical driving force. The extracellular saline contained 90 nM TTX and intracellular pipette solution contained 20 mM TEA to block sodium and potassium conductances and the spikes they facilitate. Discrete depolarisations (arrows) and the tone-evoked transduction currents are reduced in noise-exposed auditory neurons (red) compared to control (black). **E.** The amplitude of discrete depolarisations is reduced for auditory neurons from noise-exposed ears (LM t_(142)_=6.524, p<0.0001, effect size d=1.087299). The maximum six discrete depolarisations were averaged for each locust during the −100 mV voltage clamp. **F.** Example showing calculation of the latency of the tone-evoked transduction current. **G.** Latency of the transduction current was delayed for noise-exposed auditory neurons at 110 dB SPL and **H.** across sound intensities (LMEM t_(109.38)_=2.225, p=0.0281, effect size d=0.54342). **I.** The transduction current increased with louder sound amplitudes, which was well fitted with a Boltzmann equation (solid lines, Control R^2^=0.990, Noise-exposed R^2^=0.996). The transduction current was reduced for auditory neurons from noise-exposed ears (LMEM t_(138.93)_=-13.92, p<0.0001, effect size d=0.69729). Mean is plotted as circles, positive standard deviation is plotted as shaded area, transduction current amplitude from individual auditory neurons are plotted as thin shaded lines. (Control: n=12, N=6; Noise-exposed: n=12, N=7). **J.** The maximal transduction current was significantly reduced at the maximal sound intensity of 110 dB SPL.

**Figure 4.**
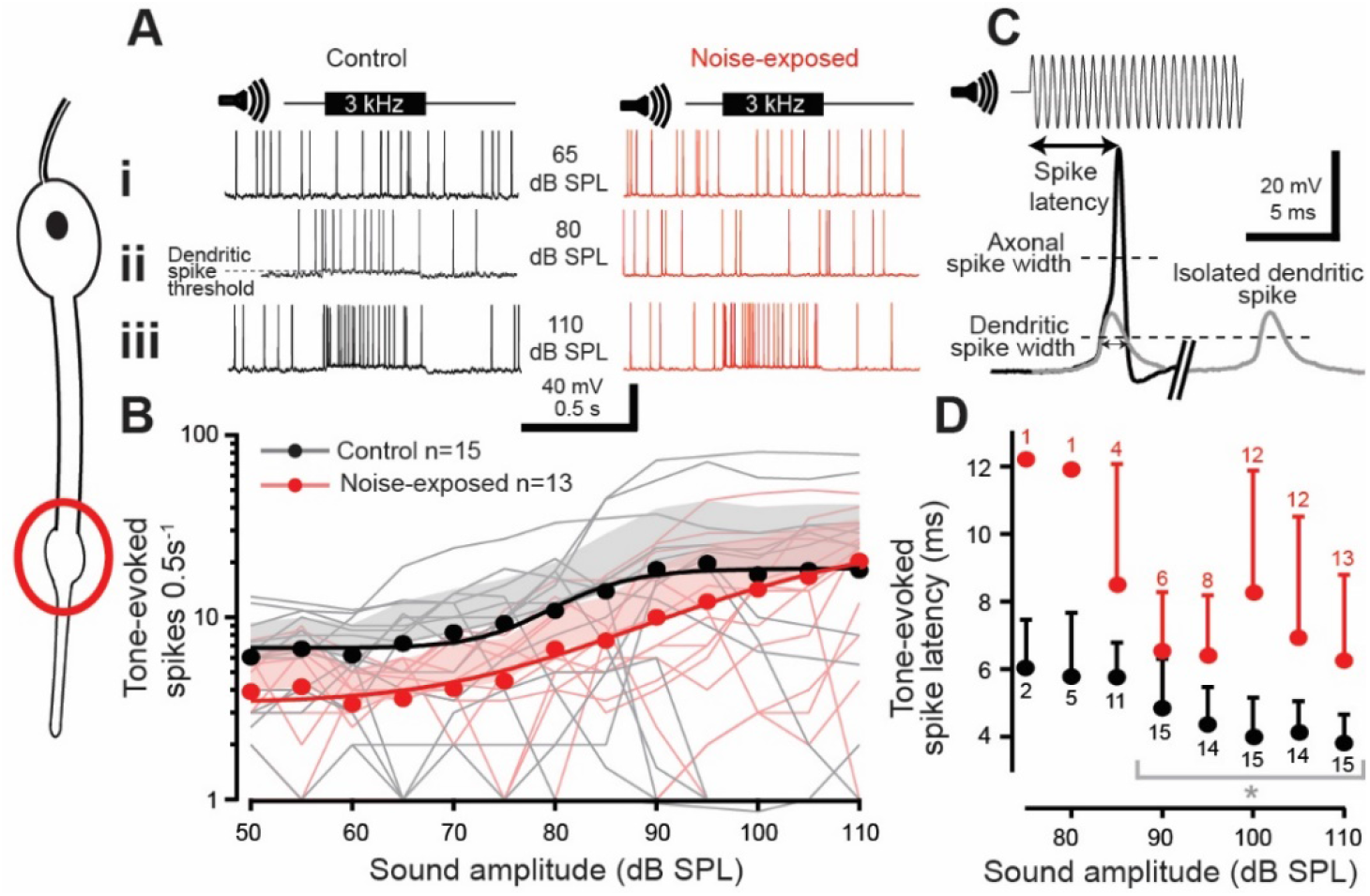
Tone-evoked spikes were recorded in current clamp mode with whole-cell patch-clamp recordings from auditory neurons from control and noise-exposed locust ears. Dendritic spikes are elicited in the apical part of the dendrite, with its presumed site highlighted by the red circle, left. **Ai, ii, iii.** Example recordings from control and noise-exposed auditory neurons when played a 0.5 s 3 kHz tone at 65, 85 and 110 dB SPL. The grey dotted line in **Aii** indicates the potential at which a dendritic spike threshold was calculated. **B.** The number of tone-evoked spikes was higher for louder sound amplitudes (LMEM t_(333.0)_=7.070, p<0.0001) which was well fitted by a Boltzmann equation (solid lines (Control: R^2^=0.965; Noise-exposed: R^2^=0.992). Tone-evoked spikes were not different for noise-exposed locust auditory neurons (red) compared to control (black) (LMEM t_(169.38)_=0.975, p=0.3309). Mean is plotted as circles, positive standard deviation is plotted as shaded area, spike counts from individual locusts are plotted as thin shaded lines (Control: n=15, N=8; noise-exposed: n=13, N=12). **C.** Example recording showing how the latency of tone-evoked spike was measured and a rare dendritic spike (grey) later in the same recording, which is overlaid on the larger axonal spike. Dotted lines indicate where the axonal spike half-width and dendritic spike half-width were measured. **D.** The latency to first spike was slower for auditory neurons from noise-exposed locusts (red) compared to controls (black) across sound amplitudes (LM t_(122)_=7.154, p<0.0001, effect size d=1.30566). Means are plotted as circles, positive standard deviation is plotted as error bars. Grey brackets with asterisk donate the recordings that were statistically tested (see Statistical Experimental design and statistical analysis for justification).

**Table 1.** There were no significant differences between the morphological or electrophysiological properties of noise-exposed and control auditory neurons (LM, p= 0.464, 0.651, 0.362, 0.7826, 0.164; t= 24.51, 0.462, −0.935, −0.280, 1.441, respectively). These measurements were recorded from spiking neurons with no TTX and TEA in the extracellular or intracellular saline. Only neurons with a resting membrane potential at least −50mV were used to compare resting potentials.

| | Dendrite length ($\mu\text{m}$) | Dendrite diameter at midpoint ($\mu\text{m}$) | Resting potential (mV) | Membrane resistance ($\text{M}\Omega$) | Capacitance (pF) |
| --- | --- | --- | --- | --- | --- |
| **Control** | 114 ± 13 (n=8) | 3.2 ± 0.74 (n=8) | 60 ± 8 (n=11) | 50 ± 47 (n=12) | 25 ± 6 (n=12) |
| **Noise-exposed** | 120 ± 15 (n=8) | 3.1 ± 0.60 (n=8) | 57 ± 5 (n=9) | 54 ± 45 (n=12) | 22 ± 5 (n=12) |

Ben isolated and optimised the transduction current at the distal ciliated end of the auditory neuron using pharmacology, voltage protocols and the optimal sound stimulus (as detailed in Figure 3). We recorded smaller discrete depolarisations in the auditory neurons of noise-exposed locust auditory neurons compared to control (Figure 3D, 3E). Discrete depolarisations are assumed to be transient stochastic opening of mechanotransduction ion channels shown both in insects (Hill, 1983b; Warren & Matheson, 2018) and vertebrate auditory receptors (Beurg et al., 2015; Pan et al., 2013). There was a significant delay in the generation of the transduction current in noise-exposed locusts compared to their control counterparts (Figure 3F) at the maximal sound amplitude of 110 dB SPL (Figure 3G) and across sound amplitudes (Figure 3H). The transduction current increased with an increased sound amplitude (Figure 3I). This dependence was well fitted with a Boltzmann equation with a clear reduction in the amplitude of the transduction current for noise-exposed locust auditory neurons. The largest difference was at their maximal transduction currents which were 146 ± 45 pA and 78 ± 25 pA for control and noise-exposed locusts respectively (figure 3J).

### Tone-evoked dendritic spikes

The transduction potential depolarises the cilium and adjacent distal dendrite of the auditory neuron, and if large enough, triggers a small dendritic spike that travels to the soma to trigger a larger axonal spike in the axon hillock (Warren & Matheson, 2018). Due to its morphology the assumed dendritic spike initiation site is the dendrite dilation that sits ~5 μm below the cilium (Figure 3C). To measure changes in tone-evoked dendritic spikes we used whole-cell patch-clamp recordings in current clamp mode. In the absence of sound stimulation spontaneous spikes were recorded which tended to be lower in noise-exposed auditory neurons (Control 5.8 ± 0.9 spikes per 0.5 s, n=15; Noise-exposed 3.4 ± 0.7 spikes per 0.5 s, n=13; LM t_(25.0)_=7.047, p<0.0001, Effect size d=0.667). The number of tone-evoked spikes increased for higher SPLs (Figure 4A). In response to a >60 dB SPL 3 kHz tone, tone-evoked spikes were triggered which closely followed a Boltzmann relationship with sound amplitude (Figure 4B). The number of tone-evoked spikes tended to be less for noise-exposed auditory neurons (Figure 4B). The latency to first spike for all neuron types decreased for increased SPLs (Figure 4C, D). The time to first spike was delayed for noise-exposed auditory neurons when compared to control auditory neurons across SPLs. We measured the width of spikes at two heights (Figure 4C) to measure possible changes in both the smaller dendritic spikes and the larger axonal spikes which they trigger. To measure axonal spike width we measured at half-height (Figure 4C, Axonal spike half-width).

To infer a measure of dendritic spike width we measured at half of the height of the dendritic spike (Figure 4C, Dendritic spike half-width) (the height of the dendritic spike was determined by taking the start of the maximum acceleration of the spike’s depolarisation as the dendritic spike height). The axonal and dendritic spike width was not different between noise-exposed and control locusts (Control axonal spike width 0.81 ± 0.24, n=15; Noise-exposed axonal spike width 0.74 ± 0.16, n=12; LM t_(25.0)_=0.53 p=0.601) (Control dendritic spike width 1.48 ± 0.44, n=9; Noise-exposed dendritic spike width 1.37 ± 0.23, n=8; LM t_(15.0)_=0.641, p=0.531). For each auditory neuron we measured the tone-elicited potential that produced an increase in the number of spikes above the background spontaneous spike rate – the dendritic spike threshold (dotted line Figure 4Aii). The dendritic spike threshold was similar for control and noise-exposed locust auditory neurons (Control dendritic spike threshold 67.1 ± 5.1 mV, n=14; Noise-exposed dendritic spike threshold 68.9 ± 5.9 mV, n=12, LM t_(24.0)_=0.822, p=0.419).

### Current-injected axonal spikes

To elicit axonal spikes (with their presumed spike initiation site in the axon hillock) current was injected into the soma through the patch electrode (Figure 5A). Current injected into either control or noise-exposed auditory neurons resulted in the generation of axonal spikes (Figure 5Ai, ii, iii). The spike rate varied as a power law of the injected current, with no difference between control and noise-exposed auditory neurons (Figure 5B). The spike latency (Figure 5C) decreased for increasing current injections (Figure 5D) and there was no meaningful difference in latency between control and noise-exposed auditory neurons (Figure 5D).

**Figure 5.**
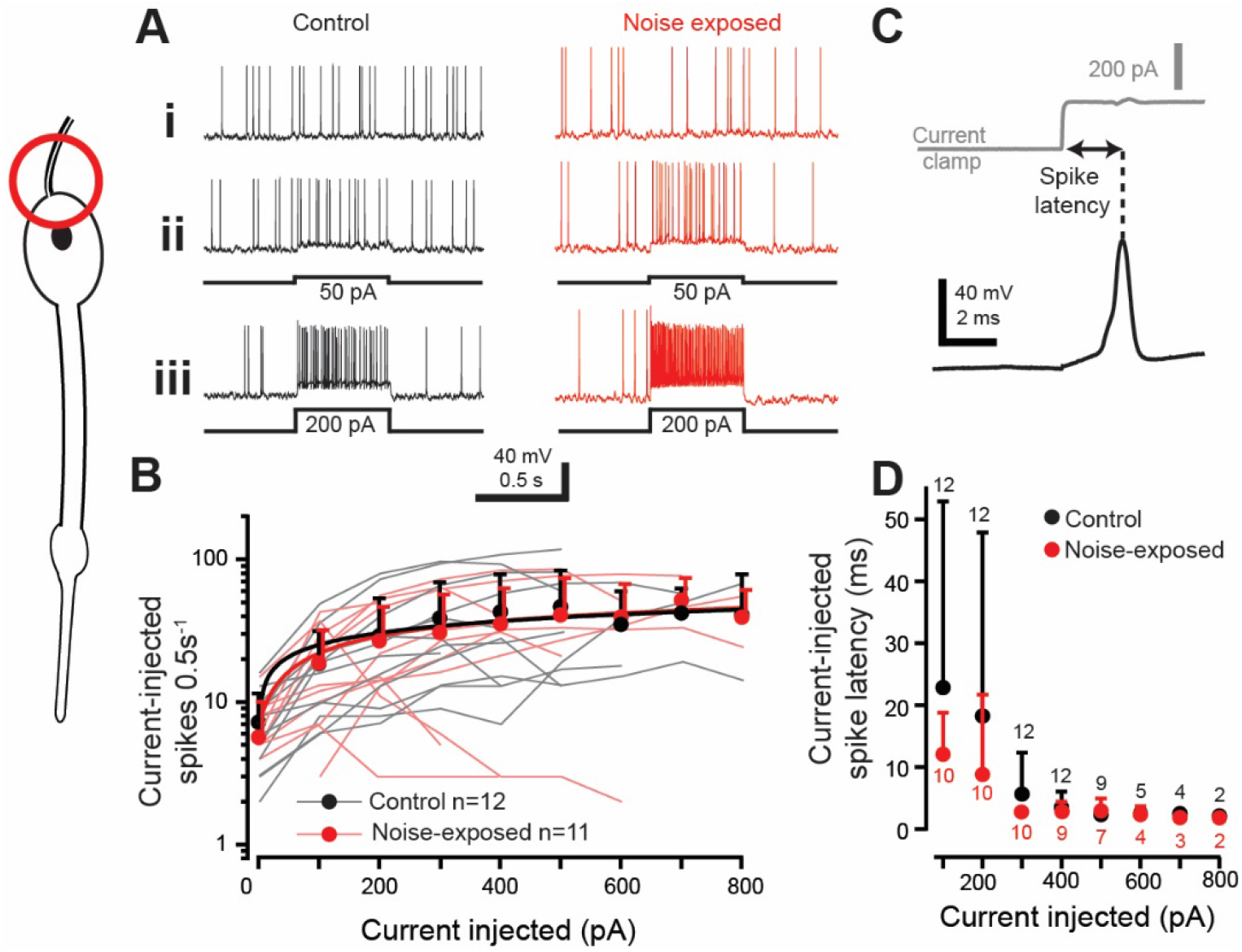
Axonal spikes were elicited through current injection into the auditory neuron somata of control and noise-exposed locust ears. Axonal spikes are assumed to be triggered in the axon hillock, red circle left. **A** Example recordings showing **Ai** spontaneous spikes and **Aii** spikes in response to 50 pA current injection and **Aiii** spikes in response to 200 pA current injection. **B.** The number of spikes triggered by current injection was fitted by a power law (solid lines) and was not different for auditory neurons from noise exposed (red) or control locusts (black) across all current injections (LMEM t_(28.38)_=-0.054, p=0.9575). Means are plotted as circles, positive standard deviation is plotted as error bars (the error bars for noise-exposed means are offset, right, for figure clarity). Current-elicited spike number from individual auditory neurons are plotted as thin shaded lines (Control: n=12, N=9; noise-exposed: n=11, N=10). **C** An example measurement of the current-injected spike latency. **D.** Current-injected spike latency across and range of injected currents, which were not different between control and noise-exposed locusts (LMEM t_(58.24)_=-1.273, p=0.208). Recordings were lost at high current injections, hence the lower n numbers at high current injections. Means are plotted as circles, positive standard deviation is plotted as error bars (One neuron from the noise-exposed group was not analysed for latency due to its high spontaneous spike rate). Grey brackets with asterisk donate the recordings that were statistically tested (see Experimental design and statistical analysis for justification).

### Streptomycin block of the mechanotransduction channels mimics noise-exposure

Ben showed a decreased amplitude of spontaneous openings (discrete depolarisations) of the mechanotransduction ion channels (Figure 3E) and decreased tone-evoked transduction current after noise exposure (Figure 3J). To mimic this effect Ben used the known mechanotransduction ion channel blocker dihydrostreptomycin to half block the mechanotransduction channels (Warren and Matheson, 2018). Blocking of the mechanotransduction ion channels had no effect on the ability to elicit spikes through current injection into the soma (Figure 6A) or their latency (Figure 6B), thus mimicking auditory neurons from noise-exposed ears. The application of 50 μM streptomycin reduced the tone-evoked spikes compared to control, which was similar to noise-exposed auditory neurons only for higher SPLs >85 dB SPL (Figure 6C). The spontaneous spike rate - which led to stark differences in the number of tone-evoked spikes at lower SPLs - was significantly lower in the presence of 50 μM streptomycin compared to control locust auditory neurons (Figure 6C) and tended to be lower compared to noise-exposed auditory neurons. The latency of tone-evoked spikes was increased in the streptomycin treated auditory neurons, compared to controls, but remained similar to the noise-exposed locusts (Figure 6D).

**Figure 6.**
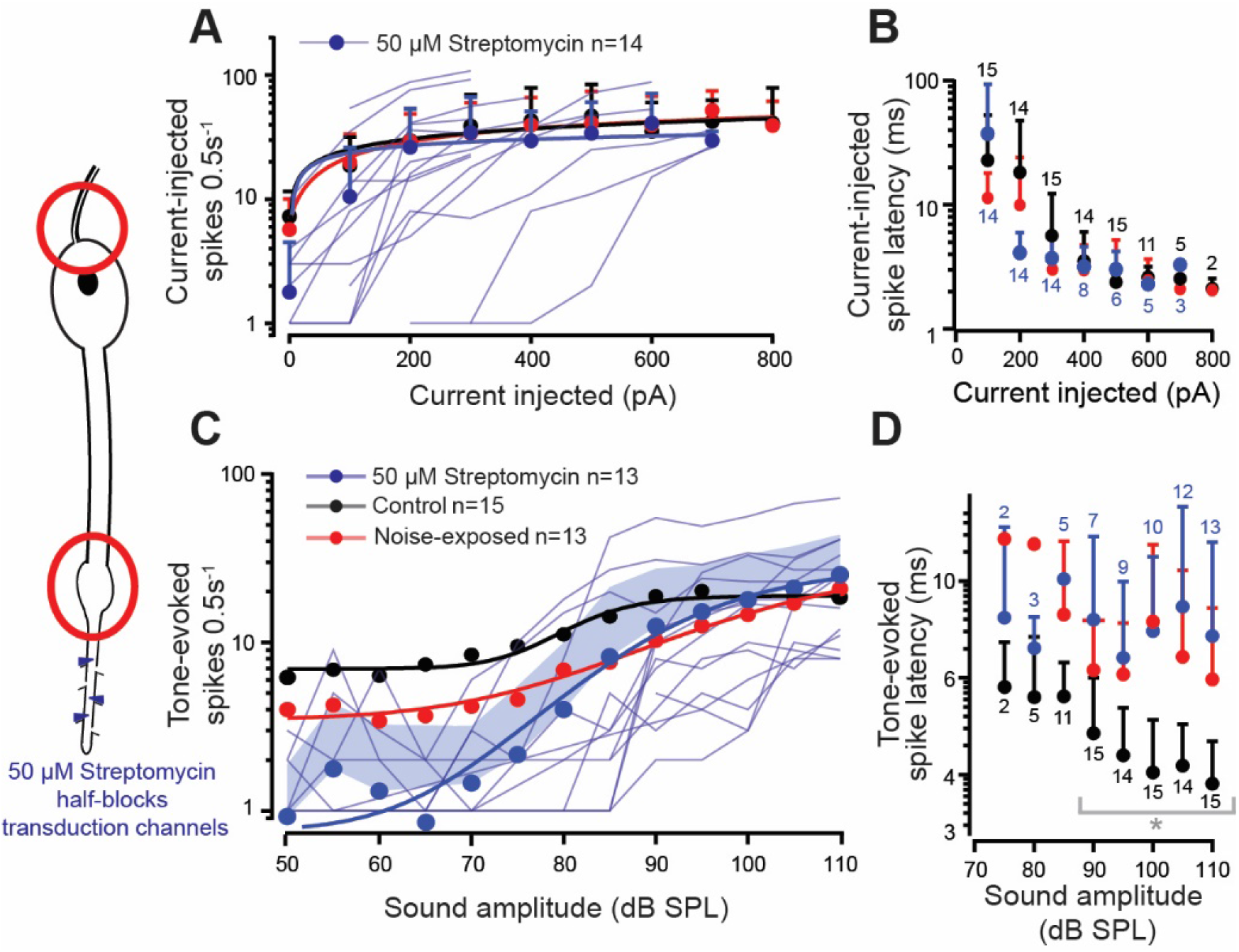
Axonal spikes at the axon hillock (red circle, top left) and dendritic spikes (red circle, bottom left) were measured in response to 50 μM streptomycin (blue hexagons, left). **A.** Number of current-injected spikes for noise-exposed (red), control (black) and streptomycin perfused (blue) auditory neurons, fitted with a power law, (solid lines) was not different between all three treatments (LMEM t_(39.38)_=0.087, p=0.9313). Means are plotted as circles, positive standard deviation is plotted as error bars, current-elicited spikes from individual auditory neurons in the presence of 50 μM streptomycin are plotted as thin blue shaded lines (Control: n=12, N=9; noise-exposed: n=11, N=10; Streptomycin perfused n=14, N=10). **B.** Latency to first current-injected spike for noise-exposed (red), control (black) and streptomycin perfused (blue) auditory neurons was not different between treatments (Control vs Noise-exposed LMEM t_(76.33)_=-1.189, p=0.237; Control vs Streptomycin LMEM t_(84.42)_=0.646, p=0.520; Noise-exposed vs Streptomycin LMEM t_(84.5)_=-1.838, p=0.0696) (plotted on logarithmic axis for figure clarity). **C.** Tone-evoked spikes for noise-exposed (red), controls (black) and streptomycin perfused auditory neurons fitted with a Boltzmann equation (Streptomycin R^2^=0.990). Streptomycin-treated auditory neurons has a lower number of tone-evoked spikes compared to controls (LMEM t_(241.76)_=-2.970, p=0.0033, similar to noise-exposed locusts’ auditory neurons (LMEM t_(244.02)_=1.869, p=0.06275). Mean is plotted as circles, positive standard deviation is plotted as shaded area and tone-evoked spike counts, from individual auditory neurons in the presence of 50 μM streptomycin, are plotted as thin blue lines. **D.** Tone-evoked spike latency for noise-exposed (red), controls (black) and streptomycin perfused (blue) auditory neurons (plotted on logarithmic axis for figure clarity). Tone-evoked spike latency was delayed in the streptomycin-treated auditory neurons compared to control (LM t_(172)_=7.241, p<0.0001), similar to auditory neurons from noise-exposed locusts (LM tt_(142)_=1.816, p=0.0711). Grey brackets with asterisk donate the recordings that were statistically tested (see Experimental design and statistical analysis for justification).

### Transcriptome analysis reveals no change in expression of the mechanotransduction ion channel candidates

The reduction in the transduction current could be explained by a reduction in expression of the mechanotransduction ion channels. To measure expression of the transduction ion channel candidates Ben Warren extracted RNA from 320 Müller’s organs from 160 control and noise-exposed locusts (two ears per locust). Andrew French analysed the RNA reads (sequenced by Beijing Genomics Institute) and quantified expression levels for the three genes that compose the two mechanotransduction ion channel candidates in insects: *nompC, nanchung* and *inactive (nanchung* and *inactive together* code a single Nanchung-Inactive ion channel) and two housekeeping genes *actin* and *GAPDH*. Andrew found no change in the expression level of any three of the genes that code the candidate mechanotransduction ion channels (Figure 7).

**Figure 7.**
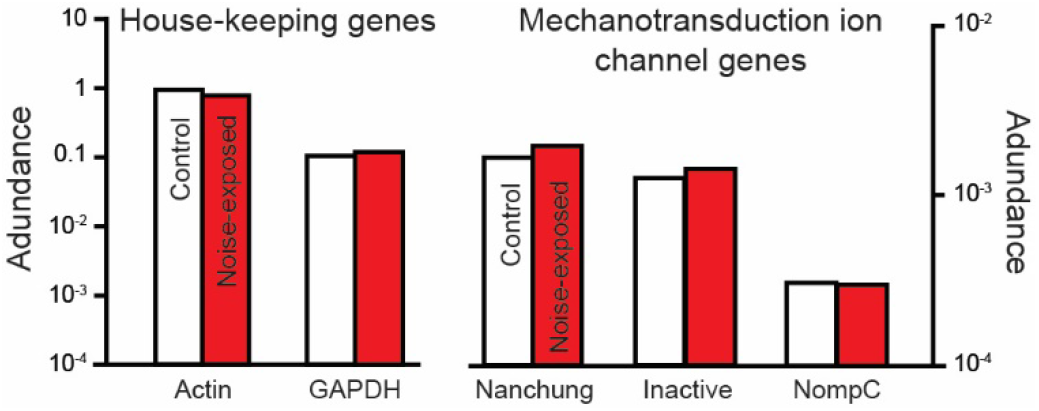
Abundance of RNA transcripts extracted from Müller’s organs of control and noise-exposed locusts for house-keeping genes, *actin* and *GAPDH* and for the three genes that together code the two mechanotransduction ion channel candidates, *nanchung, inactve* and *nompC*. Abundance was calculated by counting all matching reads to each gene’s open reading frame in the complete, groomed transcriptomes, and then normalized by the reading frame length.

## Discussion: Electrophysiological consequences in auditory neurons after noise exposure

Noise-induced hearing loss is the most common preventable sensory loss in humans. Although most noise-induced hearing loss begins with the overstimulation of primary auditory receptors we lack a characterisation of their electrical properties and electrical currents after acoustic overexposure for any animal ear. We used the physiologically accessible locust ear with its Müller’s organ attached to the inside of the tympanum to quantitatively characterise changes in mechanical vibrations of the tympanum, spikes from the auditory nerve and the electrical properties and responses of the auditory neurons themselves.

### Biomechanical responses of the tympanum

Sound-induced movements of human tympanae, and the noises they produce (otoacoustic emissions), are routinely used to assay the health of the auditory neurons buried deep within the inner ear. Thus, the tympanum is not a simple passive receiver transmitting sound-induced movements toward the auditory neurons but can also be influenced by the mechanical properties of the primary auditory receptors themselves. Although first discovered in humans (Kemp, 1978), this principle extends to the ears of reptiles (Manley et al., 2001), amphibians (Long et al., 1996), birds (Taschenberger & Manley, 1997) and multiple insects (Mhatre & Robert, 2013; Gopfert & Robert, 2003; Gopfert & Robert, 2001). We measured sound-induced movements of the locust tympanum in order to assay possible changes in the auditory Müller’s organ attached to the tympanum’s inside. We found that the tympanae of noise-exposed locust ears vibrate ~ten times higher than controls across sound levels from 50-100 dB SPL, which suggests that electro-mechanical function of Müller’s organ are altered after noise exposure. Changes in the mechanics of the locust ear will, in turn, lead to changes in the ability of the auditory neurons to transduce these mechanical movements into action potentials. Thus, we next measured the ability of noise-exposed Müller’s organs to transduce sound into action potentials that are carried along its nerve.

### In vivo electrophysiological responses of the auditory nerve

The health of an auditory system – its ability to transduce sound into electrical signals - has been assayed from the summated electrical activity of the auditory nerve (Compound Action Potentials (CAPs)) and hierarchical auditory processing areas through the brain (Auditory Brainstem Responses (ABRs)). After acoustic overstimulation there is an increase in the minimum sound level to get a sound-evoked electrical signal from the auditory nerve – the so-called auditory threshold (Coyat, et al., 2018; Christie & Eberl, 2014; Housley et al., 2013; Pilati et al., 2012; Telang et al., 2010; Sendowski et al., 2004; Ma et al., 1995; Dolan & Mills, 1989; Pettigrew et al., 1984; Van Heusden & Smoorenburg, 1981). This increased auditory threshold is mirrored by a decrease in the sound-evoked CAP amplitude after noise exposure (Christie & Eberl, 2014; Sendowski et al., 2004; Wang et al., 1992; Dolan & Mills, 1989; Pettigrew et al., 1984; Van Heusden & Smoorenburg, 1981). Due to the accessible nature of the locust’s two abdominal ears, we were able to record multiphasic nerve potentials (mainly summated action potentials) directly from the auditory nerve using hook electrodes. We found that the number of sound-elicited nerve potentials was mildly decreased and their peak response was strongly decreased in noise-exposed locust ears. In addition, the latency to first spike was significantly increased in noise-exposed locusts compared to controls which reflects increased latencies of CAPs measured in other noise-exposed auditory systems (Christie & Eberl, 2014; Sendowski et al., 2004; Pettigrew et al., 1984;; Van Heusden & Smoorenburg, 1981). Is it not known what changes take place in the auditory receptors to decrease spiking responses increase spike latency. To pinpoint noise-induced changes within primary auditory neurons we performed whole-cell patch-clamp recordings to quantify their sound-induced and spontaneous currents that lead to these auditory pathologies.

### Morphology and electrophysiological properties of the auditory neurons and the transduction current

A contributing cause of deafness is loss of auditory hair cells (in vertebrates) through necrotic and apoptotic pathways, which can take place within 48 hours of acoustic trauma (Coyat et al., 2018; Ryals et al., 1999; Cotanche, 1987; Hamernik et al., 1984). We thus, measured the morphology of recorded auditory neurons exposed to 24 hrs of loud sound (126 dB SPL) but found no difference compared to controls. The electrical properties of the auditory neurons such as their membrane resistance, resting membrane potential and capacitance was also not affected by noise exposure, although the power of our analyses was limited by our sample size.

### Physiological basis of a decrease in the transduction current

We then moved on to analyse changes in the spontaneous and sound-evoked openings of the transduction ion channels. Spontaneous channel openings, in the absence of sound, manifest as discrete depolarisations of variable magnitude, which upon auditory stimulation, summate to produce the transduction current (Hill, 1983a; Warren & Matheson, 2018). There was a significant reduction in the size of the discrete depolarisations and the magnitude of sound-evoked transduction current and an increase in the latency to elicit the transduction current. The decrease in the sound-evoked transduction current paralleled the single published example recording showing this in a vertebrate hair cell seconds after mechanical stimulation (Cody & Russell, 1985). The findings here have multiple explanations: 1 the electrochemical driving force for the ions passing through the transduction channels is decreased; 2 the mechanical attachment to the auditory neurons - and the sound-induced force delivered to them - has weakened; 3 the number of transduction channels is reduced. Ciliated auditory receptors across animals function with common biophysical principles (Albert et al., 2007; Howard & Hudspeth, 1988), share striking genetic homology (Wang et al., 2002) and (most probably) evolved from the same ancestral auditory receptor (Fritzsch & Beisel, 2004). Thus, we address each of these three explanations in a comparative context with other auditory systems across the animal kingdom.

#### Explanation 1: a decrease in the electrochemical driving force across the mechanotransduction ion channels

We have shown no large reduction in the intracellular potential of the auditory neurons, despite a large decrease in the transduction current. Thus, if a decrease in electrochemical gradient causes a decreased transduction current, it must result in a decrease in the extracellular electrochemical potential maintained in a specialised receptor lymph cavity that baths the external surface of the transduction channels. A decrease in ionic gradient and potential in the receptor lymph could result from damage in the supporting cells that pump ions. Indeed, in the fruit fly, knocked down expression of the sodium pump (Na^+^/K^+^ ATPase) in the supporting scolopale cells that enclose the receptor lymph result in deafness and anatomical pathologies in the receptor lymph space and sensitisation to acoustic trauma (Christie & Eberl, 2014; Roy et al., 2013). A similar result is found in mammals where the supporting cells (in the stria vasculairs) are oxidatively damaged (Shi et al., 2015) after noise exposure and, in mammals, there is a concurrent decrease in the endocochlear potential (Hirose & Liberman, 2003) and normal concentrations of potassium and calcium ions (Li et al., 1997; Ma et al., 1995; Vassout, 1984). Thus, there is a precedent for a decrease in the extracellular potential in the receptor lymph cavity based on comparative work in other animal ears.

#### Explanation 2: a decrease in the sound-induced force delivered to the auditory neurons

The transduction current in noise-exposed auditory neurons is reduced by half despite a ten-fold increase in sound-evoked tympanal displacements. The increased movement of the locust tympanum mirrors a likely increase in compliance of the human tympanum measured in soldiers exposed to impulse noise (Job et al., 2016). In the locust ear there must be a drastic decrease in mechanical coupling between the tympanum and the auditory neuron cilia where these sound-induced forces open transduction ion channels, otherwise we should expect an increase of the transduction current after noise exposure. Despite this, the maximal transduction current asymptotes for sound amplitudes approaching 110 dB SPL (Figure 3I) suggesting that all the transduction channels are opened at these higher sound levels. All together it seems unlikely that a reduction in the sound-induced force is the main cause of the decreased transduction current.

#### Explanation 3: a decrease in the transduction channel number

The third explanation is particularly hard to unequivocally test because the protein/s that form/s the transduction channel have, controversially, not been identified in insects – or indeed any animal ear (although in mammals, it seems, they are getting close (Qiu & Müller, 2018)). Despite this, the only ion channels linked to hearing in insects, and also expressed in the cilium, where transduction takes place, are NompC and Nanchung-Inactive (Gong et al., 2004; Kim et al., 2003). We showed no change in expression of the three genes that compose the two candidate mechanotransduction ion channels. Providing that either Nanchung-Inactive or NompC are the transduction channel, the decrease in the transduction current is probably due to a decreased electrochemical potential in the receptor lymph cavity that bathes extracellular surface of the, still elusive, mechanotransduction ion channel.

### Pathologies resulting from a decreased transduction current

A decreased transduction current results in fewer sound-evoked spikes for all but the loudest sound intensities and delayed sound-evoked spikes. We tested if potential pathologies in the spike generating machinery in the axon hillock resulted in fewer sound-evoked spikes but found no difference in the threshold or latency between noise-exposed and control auditory neurons. We found no changes in the dendritic spike threshold or their spike width and no difference in the axonal spike width, the decrease in auditory performance, at least in an insect ear, is due to a decreased transduction current. To test this hypothesis we used a known mechanotransduction ion channel blocker dihydrostreptomycin (Marcotti et al., 2005; Warren & Matheson, 2018) at a concentration to mimic the decrease in transduction current due to noise exposure. We found no change in the spiking threshold or latency to first spike upon current injection to the soma but sound-evoked spikes were decreased and delayed similar to the noise-exposed auditory neurons. The decreased tone-evoked dendritic spikes is presumably a direct consequence of the reduced transduction current.

### Conclusions

This study presented here is the first systematic assessment of changes in primary auditory receptors in any animal after noise-exposure. We found changes in the transduction current magnitude that translated into decreased sound-evoked spikes. A beneficial consequence of the reduced transduction current are the reduced metabolic demands put on an auditory system, which may preserve its function. It reduces the metabolic demand through two processes: 1. It reduces the sound-evoked transduction current that flows through the neurons and which the supporting scolopale cell continually expend energy, through transmembrane pumps, to recycle these ions to the auditory neurons apical end. 2. It reduces the sound-evoked spikes in the auditory neurons themselves thus sparing the metabolically demanding process of pumping sodium and potassium ions out and in to the auditory neuron to maintain spike readiness. This is an innovative, at least short-term, solution to protect against further damage by auditory overstimulation – to simply reduce the incoming signal at its source. In mammals the shown decrease in the electrochemical driving force, across hair cells (Hirose & Liberman, 2003), would, in theory, lead to a decreased transduction current – similar to that measured here in the locust ear. Thus, we predict the transduction potential in hair cells is also reduced which results in a similar protective mechanism across auditory systems including mammalian hair cells. A comparative analysis of hearing across animals is a powerful approach to understand the basic tenets of auditory transduction. The strengths of each model system can be exploited to give us deeper insights into common principles of how animals hear (Albert et al., 2007; Palghat et al., 2012) and our analysis here has special relevance for understanding the most preventable sensory impairment in humans, noise-induced hearing loss.

## Materials and methods

### Locust husbandry

Desert locusts (*Schistocerca gregaria*) were reared in crowded conditions (phase gregaria) on a 12:12 h light:dark cycle at 36:25°C. Locusts were fed on a combination of fresh wheat and bran *ab libitum*. Male locusts between 10 and 20 days post imaginal moult were used for all experiments. We are willing and open to share our locust strains with other research groups.

### Noise conditioning and acoustic stimulation

The wings of all locusts were cut off at their base to increase the exposure of the conditioning tone to their tympanal ears, which are otherwise covered by their wings. Up to twenty locusts, for both the noise-exposed group and the control group, were placed in a cylindrical wire mesh cage (8 cm diameter, 11 cm height). Both cages were placed directly under a speaker (Visaton FR 10 HM 4 OHM, RS Components Ltd). For the noise-exposed group only, the speaker was driven by a function generator (Thurlby Thandar Instruments TG550, RS Components Ltd) and a sound amplifier (Monacor PA-702, Insight Direct Ltd) to produce a 3 kHz tone at 126 dB SPL, measured at the top of the cage where locusts tended to accumulate. Sound Pressure Levels (SPLs) were measured with a microphone (Pre-03 Audiomatica, DBS Audio) and amplifier (Clio Pre 01 preamp, DBS Audio). The microphone was calibrated with a B&K Sound Level Calibrator (CAL73, Mouser Electronics). The locust ear was stimulated with the same speaker and amplifier as above for hook electrode recordings. For intracellular recordings from individual auditory neurons the speaker was driven by a custom made amplifier controlled by a EPC10-USB patch-clamp amplifier (HEKA-Elektronik) controlled by the program Patchmaster (version 2×90.2, HEKA-Elektronik) running under Microsoft Windows (version 7).

### Biomechanical measurements of the tympanum with laser Doppler vibrometry

For *in vivo* measurements the locusts wings and hind legs were cut off and locusts were fixed so that their tympanum was perpendicular to the micro-scanning Laser Doppler Vibrometer (PSV 300, Polytec, Waldbronn, Germany) with a close up unit (OFV 056). A loudspeaker (ESS Air Motion Transformer, South El Monte, CA, USA) was placed at least 10 cm away, to avoid operating in the near field. A microphone (Bruel & Kjaer 4138, Naerum, Denmark) was positioned to measure the sound pressure at the tympanal membrane.

For *ex vivo* measurements whole ears, including Müller’s Organ attached to the internal side of the tympanum, were dissected from the first abdominal segment, by cutting around the small rim of cuticle surrounding the tympanum with a fine razor blade. Trachea and the auditory nerve (Nerve 6) were cut with fine scissors (5200-00, Fine Science Tools), and the trachea and connective tissue removed with fine forceps. The ear was secured, inner side up, into a 2 mm diameter hole in a Perspex divider (go to www2.le.ac.uk/departments/npb/people/bw120 to download the locust ear holder file for 3D printing) using an insect pin pushed through the anterior rim of cuticle and into 2 mm of Sylgard (184 Silicone Elastomer, Dow Corning) on the base of a 30 mm diameter petri dish. The tympanum in its holder was positioned at an angle of 30° off vertical to observe Group-III neurons of Müller’s Organ from above. A watertight seal was made between the ear cuticle and the divider hole with dental glue (Protemp 4, 3M ESPE) and nerve 6 was secured into the glue at the ventral border of the tympanum.

### In vivo hook electrode recordings from auditory nerve six

Locusts were secured ventral side up in plasticine. A section of the second and third ventral thoracic segment was cut with a fine razor blade and removed with fine forceps. Tracheal air sacks were removed to expose nerve six and the metathoracic ganglia. Hook electrodes constructed from silver wire 18 μm diameter (AG549311, Advent Research Materials Ltd) were hooked under the nerve and the nerve was lifted out of the haemolymph. Signals were amplified 10,000 times by a differential amplifier (Neurolog System) then filtered with a 500 Hz high pass filter and a 50 kHz low pass filter. This amplified and filtered data was sampled at 25 kHz by Spike2 (version 8) software running on Windows (version 10).

### Dissection of Müller’s Organ and isolation of Group-III auditory neurons

For intracellular patch-clamp recordings from individual auditory neurons the abdominal ear was excised and placed into a preparation dish as explained above (Biomechanical measurements of the tympanum with laser Doppler vibrometry). This preparation allowed perfusion of saline to the internal side of the tympanum, necessary for water-immersion optics for visualizing Müller’s Organ and the auditory neurons to be patch-clamped, and concurrent acoustic stimulation to the dry external side of the tympanum. The inside of the tympanum including Müller’s Organ was constantly perfused in extracellular saline.

To expose Group-III auditory neurons for patch-clamp recordings, a solution of collagenase (0.5 mg/ml) and hyaluronidase (0.5 mg/ml) (C5138, H2126, Sigma Aldrich) in extracellular saline was applied onto the medial-dorsal border of Müller’s Organ through a wide (12 μm) patch pipette to digest the capsule enclosing Müller’s Organ and the Schwann cells surrounding the auditory neurons. Gentle suction was used through the same pipette to remove the softened material and expose the membrane of Group-III auditory neurons. The somata were visualized with an upright microscope (BH-2, Olympus) using a water immersion objective (W Plan-APOCHROMAT, 40x, 1.0 numerical aperture, 2.5 mm working distance, Zeiss) and differential interference contrast optics.

### Electrophysiological recordings and isolation of the transduction current

Electrodes with tip resistances between 3 and 4 MΩ were fashioned from borosilicate class (0.86 mm inner diameter, 1.5 mm outer diameter; GB150-8P, Science Products GmbH) with a vertical pipette puller (PP-830, Narishige). Recording pipettes were filled with intracellular saline containing the following (in mM): 190 K-aspartate, 4 NaCl, 2 MgCl2, 1 CaCl2, 10 HEPES, 10 EGTA. To block K^+^ channels necessary for isolation the transduction current 20 mM tetraethylammonium chloride (TEA) was added to the intracellular saline, K-aspartate was reduced to 170 mM to maintain the same osmolality. To isolate the transduction current we also blocked spikes with 90 nM Tetrodotoxin (TTX) in the extracellular saline. During experiments, Müller’s Organs were perfused constantly with extracellular saline containing the following in mM: 185 NaCl, 10 KCl, 2 MgCl2, 2 CaCl2, 10 HEPES, 10 Trehalose, 10 Glucose. The saline was adjusted to pH 7.2 using NaOH. The osmolality of the intracellular and extracellular salines’ were 417 and 432 mOsm, respectively.

Dihydrostreptomycin sesquisulfate, 50 μM (D7253, Sigma Aldrich) was used to block mechanotransduction ion channels. Dihydrostreptomycin sesquisulfate was perfused at least 15 min before recordings. Whole-cell voltage-clamp recordings were performed with an EPC10-USB patch-clamp amplifier (HEKA-Elektronik) controlled by the program Patchmaster (version 2×90.2, HEKA-Elektronik) running under Microsoft Windows (version 7). Electrophysiological data were sampled at 50 kHz. Voltage-clamp recordings were low-pass filtered at 2.9 kHz with a four-pole Bessel filter. Compensation of the offset potential and capacitive current were performed using the “automatic mode” of the EPC10 amplifier. The calculated liquid junction potential between the intracellular and extracellular solutions was also compensated (15.6 mV for normal saline and 13.5 mV for TTX and TEA saline; calculated with Patcher’s-PowerTools plug-in from www3.mpibpc.mpg.de/groups/neher/index.php?page=software). Series resistance was compensated between 50 and 70% with a time constant of 100 μs.

### Staining and confocal microscopy

To stain Group-III auditory neurons, recording electrodes were filled with Neurobiotin (1% m/v, SP-1120, Vector Laboratories) dissolved in intracellular saline. To aid diffusion of Neurobiotin into the neurons a positive current of ~200 pA was injected for ~30 min. Directly after staining, Müller’s Organs were fixed overnight at 5°C in 4% paraformaldehyde (P6148, Sigma Aldrich) dissolved in Phosphate Buffer Saline (PBS). Müller’s Organs were then washed three times in PBS then gently shaken at room temperature for 20 min in a mixture of collagenase (0.5 mg/ml) and hyaluronidase (0.5 mg/ml) (C5138 and H2126, Sigma Aldrich). They were washed three times in PBS (3×10 min) then gently shaken at room temperature in 0.2 % m/v Triton-X100 dissolved in PBS (2×60 min). Müller’s Organs were then gently shaken in 20 μg/ml Dylight 488 strepavidin (SA-5488, Vector Laboratories) and 0.05 mg/ml DAPI (D9542, Sigma Aldrich) in PBS overnight at 5°C, washed three times in PBS (3×10 min), dehydrated in an ethanol series and cleared in Methyl salicylate (M6752, Sigma Aldrich).

Fluorescence images (pixel size 0.31 μm2, z stacks of 0.31 μm) were captured with a confocal microscope (FV1000 CLSM, Olympus) equipped with Plan-UPlanSApo 10x (0.4 numerical aperture) and 20x (0.75 numerical aperture) lenses. Fluorescence emission of Dylight 488 was collected through a 505-530 nm bandpass filter. Confocal images were adjusted for contrast and brightness, overlaid and stacked in ImageJ (version 1.51, National Institutes of Health). The ImageJ plugin Simpler Neurite Tracer was used to determine the distance from the centre of the soma to the dendrite dilation (Fig. 3C).

### RNA extraction, sequencing and transcriptome analysis

A total of 320 Müller’s organs from 160 control locusts (two ears per locust) and 320 Müller’s organs from 160 noise-exposed locusts were extracted by grasping the Müller’s organ through the tympanum with fine forceps and pulling it out. Müller’s organs were placed in an Eppendorf tube submerged in liquid nitrogen. RNA was extracted and then treated with DNase using a Thermo-fisher RNAqueous kit (AM1931, ThermoFisher). RNA was shipped in dry ice to Beijing Genomics Institute (Hong Kong) for sequencing. The samples were quality checked and had RNA Integrity Values (RIN) of 8.7 and 8.4 for control and noise-exposed samples, respectively. The two samples were sequenced using Illumina HiSeq 2000 using a 100 paired ends module. We obtained 186.1 million clean reads for RNA extracted from the Müller’s organs of noise-exposed locusts and 185.9 million clean reads from the Müller’s organs of control locusts. Andrew French analysed the RNA reads and quantified expression levels (abundances) for the three genes that compose the two mechanotransduction ion channel candidates in insects: nompC, nanchung and inactive (nanchung and inactive together code a single heteromeric Nanchung-Inactive ion channel) and the two housekeeping genes actin and GAPDH. Initial cDNA reads were groomed to have 80 or more contiguous nucleotides with Phred score >19 to give a final database of ~100 million pairs of reads. Sequences of interest were identified by searching all possible translations of reads from the control transcriptome versus amino acid sequences of published insect genes, including *Drosophila melanogaster*, using BLOSUM matching matrices (Henikoff & Henikoff, 1993). Identified reads of interest were extended by the transcriptome walking algorithm (French, 2012) using an initial minimum overlap of 60 nucleotides. Walking was always continued to identify the complete protein coding sequence, including both START and STOP codons.

Relative abundances of transcribed mRNA sequences in the two tissues were estimated by searching each complete groomed transcriptome library for reads matching the reading frame of each gene, using the criterion of at least 90/100 identical nucleotide matches to score each read as derived from the gene. Matching reads as a fraction of total reads counted were then normalized by gene length and expressed as abundance relative to the most abundant actin gene in the control tissue.

### Experimental design and statistical analysis

For all recordings we used male *Schistocerca gregaria* from the Leicester ‘Locust Labs’ laboratory stock. Throughout the manuscript n refers to the number of recorded neurons and N refers to the number of Müller’s Organ preparations used to achieve these recordings (i.e. n=10, N=6 means that 10 neurons were recorded from 6 Müller’s Organs). All n numbers are displayed on the figures for clarity. The Spread of the data is indicated by 1 standard deviation as the standard deviation indicates the spread of the data. Median and Q1 and Q3 are displayed by bars when individual measurements are plotted. The treatment of the locust: noise-exposed or control, remained blinded to the experimenter until data analysis was completed. To test for differences and interactions between control, noise-exposed and streptomycin treated locusts we used either a linear model (LM) or Linear Mixed Effects Model (LMEM) when repeated measured are reported. Models were fitted in R (Version 3.4.3). The test statistic for these analyses (t) are reported with the degrees of freedom (in subscript) and p value, which are approximated using Satterthwaite equation. We report Cohen’s d effect size for significant differences. Curves where fitted to the data using Matlab (version R2018a). The shape of the data was described by fitting Boltzmann voltage equation using non-linear least squares method. For each curve fit the goodness of fit is indicated by R^2^. For testing differences between the latency of tone-evoked spikes (Figure 4D and 6D) we analysed the highest five SPLs because: 1. the higher n numbers at these louder SPLs provided more rigorous power, 2. The latencies at these higher SPLs reached a plateau and therefore did not markedly change, and 3.

This is the first study to measure the effects of noise exposure on the locust’s auditory system. Thus, we had no *a priori* effect size for power calculations but we will able to use the effect sizes reported here for power calculations for future studies using the locust ear as a model for hearing loss.

## Acknowledgements

BW was funded by the Royal Society and the Leverhulme Trust. Within the University of Leicester BW was was also funded by the Department of Neuroscience, Psychology and Behaviour and the Wellcome Trust Institutional Strategic Support Fund. GF was funded by a Royal Society Enhancement Award. Work carried out by EK and JFCW was funded by the European Research Council under the European Union’s Seventh Framework Programme (FP/2007-2013) / ERC Grant Agreement n. 615030. Andrew French is funded by Discovery Grant RGPIN/03712 from the Natural Sciences and Engineering Research Council of Canada. We would also like to thank Neil Rimmer and Jake Cranston for locust husbandry and Ben Cooper and Brendan O’Connor for help with statistical analysis and making sure the locust treatment remained blinded to the experimenter. I would also like to thank Nathan Suray for performing preliminary experiments for *in vivo* hook electrode recordings.

## Author contributions

BW conceived the idea for the paper, collected and analysed the patch-clamp electrophysiological data, composed Figures 3–6 and wrote the paper. GF collected all data for the *in vivo* hook electrode recordings, analysed the data and composed Figure 1 and performed all statistical analyses. EK collected and analysed the mechanical laser Doppler data and composed Figure 2. AF analysed and interpreted the transcriptome data and composed Figure 7. GF, JCMW, EK and AF helped refine figures, write the paper and interpret data.

## Competing Interests

The authors declare no competing interests.

